# Isolation and characterisation of novel fruit bat alphaherpesvirus from *Rousettus aegyptiacus* bats in Coastal Kenya

**DOI:** 10.64898/2026.06.25.734443

**Authors:** George Kisoi, Joel Bargul, Johnson Kinyua, Solomon Langat, Hellen Koka, Joel Lutomiah, Fredrick Eyase

## Abstract

**Background:** Herpesviruses are a group of double-stranded DNA viruses known to infect a wide range of vertebrates and establish life-long latent infections. While bats serve as natural reservoir hosts for numerous viral families, relatively few bat herpesviruses have been successfully isolated. In this study, we report the isolation and characterization of two novel alphaherpesvirus strains obtained from *Rousettus aegyptiacus* bats in Coastal Kenya.

**Methods:** The samples of oral and rectal swabs were collected from three different species of bats from coastal Kenya between October 2024 and April 2025; the bat species collected include *Hipposideros* spp., *Coleura afra*, and *Rousettus aegyptiacus*. Virus isolation was performed by inoculation of samples in Vero E6 cells and subsequent monitoring for cytopathic effects (CPE). Total nucleic acids were extracted from CPE positive cultures and subjected to library preparation to enable unbiased detection of both RNA and DNA viruses. The libraries were sequenced using next-generation sequencing with Illumina MiSeq platform. Subsequently, bioinformatic analysis was carried out to identify the virus, generate consensus genomes as well as phylogenetic analysis to determine the placement of identified viruses.

**Results:** Two samples from *R. aegyptiacus* (KIK_460_O and KIK_465_O) induced typical CPE within five days. Sequencing and assembly yielded partial consensus sequences of approximately 60 kb (KIK_460_O) and 70 kb (KIK_465_O), representing extended genomic data for a bat-associated alphaherpesvirus. This virus has a genome of about 140kb, indicating that our partial assemblies account for about 43-50% of the total genome. Both isolates were found to be closely related to Dzifa herpesvirus, an alphaherpesvirus previously identified in Kilifi, Kenya. Alphaherpesvirus was identified based on partial sequencing of *UL19* (3,787bp) and *UL30* (2,846bp) genes. The two isolates were found to be identical at the *UL19* gene, showing that they belonged to the same virus strain. Phylogenetic analysis showed that the novel alphaherpesvirus belongs to primate alphaherpesviruses under the subfamily *Alphaherpesvirinae*.

**Conclusion:** This study reports the isolation and genomic characterization of a novel fruit bat alphaherpesvirus from Kenyan *Rousettus aegyptiacus* bats. The partial genome assembly (60-70 kb) represent the first extended genomic data for this virus, covering approximately 43-50% of the estimated 140 kb complete genome. The phylogenetic placement of this alphaherpesvirus near primate viruses, especially Pteropodid alphaherpesvirus 1, suggests bat-association and needs further investigation into its zoonotic potential.

## 1. Introduction

Herpesviruses are globally distributed double stranded deoxyribonucleic acid (dsDNA) viruses that infect a diverse array of vertebrate species, ranging from mammals, birds, reptiles, to fish [1]. They are unique among dsDNA viruses for their capacity to infect their natural host lifelong in a latent state, with periodic reactivation phases that assist in the sustainability and transmission of the virus [2]. The *Herpesviridae* family includes three subfamilies of viruses depending on their genome structure, life cycle, and tropism, which are the *Alphaherpesvirinae*, *Betaherpesvirinae*, and *Gammaherpesvirinae* [3]. Although they are known for their high level of host specificity due to their long history of co-evolution with the host, herpesviruses can occasionally cross between species, resulting in severe consequences for the latter as is the case for *Macacine alphaherpesvirus 1* (B virus) in humans [4]. Among bats, numerous viruses have been discovered, including herpesviruses [5]. However, despite the global distribution and ecological importance of bats, relatively few bat herpesviruses have been isolated and fully characterized [6]. The most detailed study of a bat alphaherpesvirus up to date is the identification of fruit bat alphaherpesvirus 1 found in Pteropus bats from Indonesia [7].

Bats are mammals belonging to the order of Chiroptera. They are made up of over 1,500 different species and act as important reservoir hosts for many viruses [8]. Bats in Africa, especially in East Africa, are very diverse and abundant, inhabiting a variety of ecological niches, such as from savannahs to tropical forests and caves [9]. Their wide distribution facilitates the maintenance and transmission of diverse viral communities. They are widely acknowledged as important elements of the world ecosystem owing to their contributions in regulating insects, pollination, and dispersal of seeds [10]. In addition to these ecological capabilities, they have received considerable scientific interest as hosts of a large range of viruses, some of which have high zoonotic potential [11]. In African cave systems, bats often occupy ecological niches, such as forest canopies, and fragmented habitats that allow roosting, foraging, and nesting activities that provide chances of exchanging viruses [12,13]. This ecological overlap has facilitated historical cross-species transmission events, including spillover of RNA viruses such as Ebola and Marburg between bats and humans [14].

The ecological overlap between bats, primates, and humans in shared habitats, such as the limestone caves and coastal forests of East Africa, may facilitate cross-species viral transmission [15,16]. Recent metagenomic studies have revealed that bats harbor herpesviruses which are phylogenetically related to those found in primates, suggesting either ancient host-switching or a shared co-evolutionary history [17,18].

While RNA viruses such as filoviruses, coronaviruses, and paramyxoviruses have been extensively investigated in bats because of their epidemic potential, bat-associated DNA viruses remain comparatively understudied [17,19]. Among these, herpesviruses are of particular interest from a One Health perspective due to their ability to establish lifelong latent infections, reactivate under conditions of immunosuppression, and undergo recombination during co-infections, potentially generating variants with altered host range and virulence [20,21]. Moreover, the long-term co-evolution of herpesviruses with their hosts provides valuable insights into virus–host relationships and historical host-switching events [22,23]. Given the ecological overlap between bats, non-human primates, and humans in East Africa, and the limited characterization of bat herpesviruses in the region, investigating these viruses is important for understanding viral diversity, evolution, and emergence potential [24].

Therefore, this study aimed to isolate and characterize viruses from bats inhabiting the coastal regions of Kenya and to assess their evolutionary relationships with known herpesviruses from other host species.

## Materials and Methods

### Ethical considerations

Scientific and ethics approval was obtained from the Scientific and Ethics Review Unit (SERU) of the Kenya Medical Research Institute (KEMRI) under Protocol No. KEMRI/SERU/CVR/002/4729. The National Commission of Science, Technology, and Innovation (NACOSTI) granted the research authorization under license No. NACOSTI/P/23/27916.

### Sample collection

Between October, 2024 and April 2025, samples were collected at major chiropteran roosting sites across the coastal regions of Kilifi and Kwale Counties, Kenya (Figure 1), an area selected for its coastal forest fragments, limestone caves, and fruiting vegetation [25]. In Kwale County, sampling focused on the main entrance of the Shimoni cave system. Within Kilifi County, sampling was done in Malindi, a coral forest of the Kikambala area, and roosts within an abandoned quarry in the Mavueni area. The three predominant bats included: *Hipposideros* spp. (n = 50), *Coleura afra* (n = 50) and *Rousettus aegyptiacus* (n = 70). Bats were swabbed using sterile polyester-tipped swabs wetted with anti-fungal and antibiotics solutions (Amphotericin B 0.25 μg/mL, Streptomycin 100 μg/mL) were used. To guarantee sufficient mucosal contact, swabs were carefully placed into the rectum and oral cavity and rotated. A total of 170 oral and rectal swabs were collected [26,27]. Following sampling, each bat was promptly released after being morphologically identified in the field using taxonomic keys [28]. Samples were put into cryovials containing 2.0 mL of Viral Transport Media (VTM) and stored in liquid nitrogen and shipped on dry ice to Kenya Medical Research Institute (KEMRI) biosafety level 3 (BSL-3) virology lab for processing.

**Figure 1:**
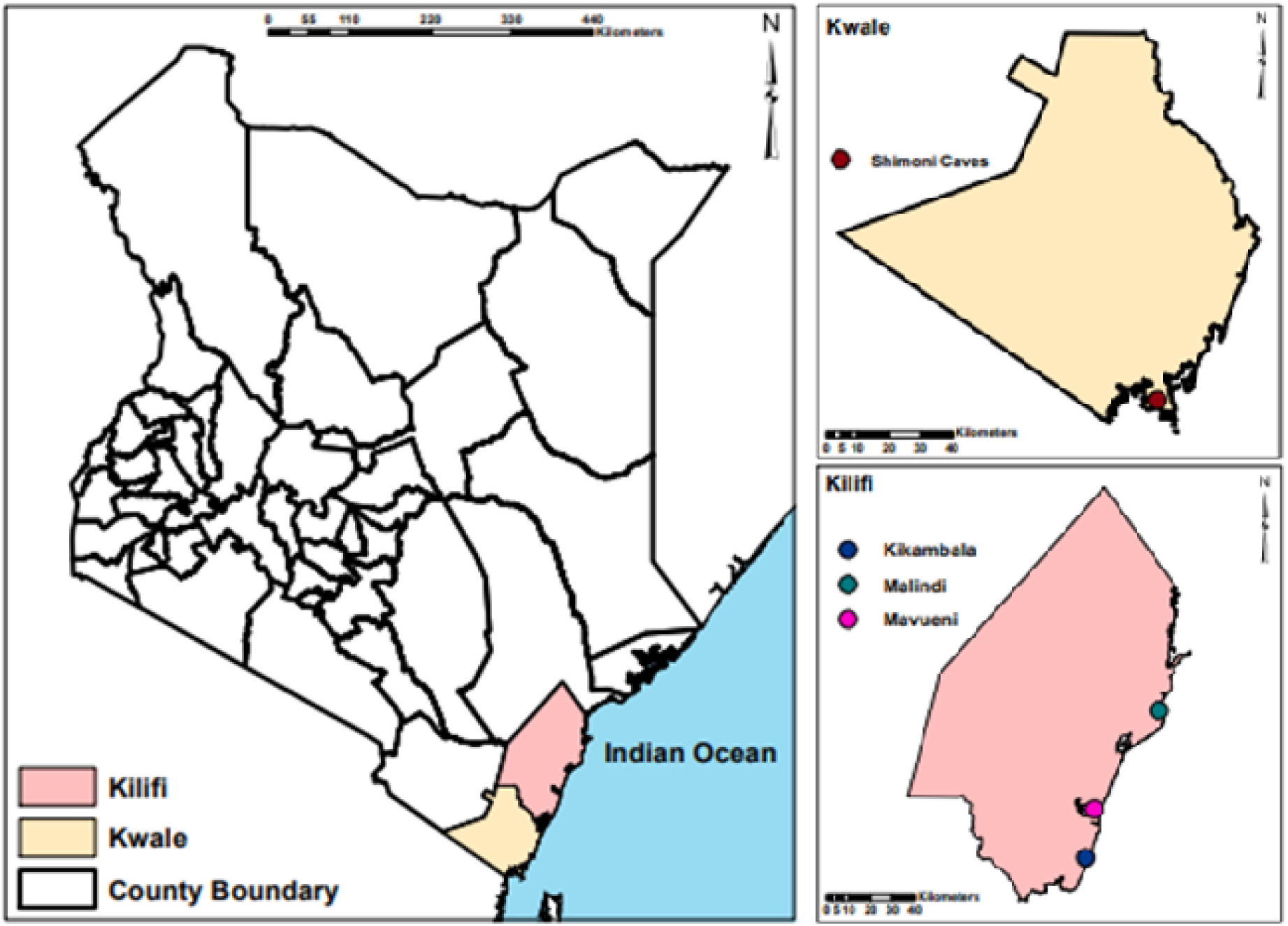
Map of coastal Kenya showing the sampling sites in Kilifi and Kwale Counties.

The map highlights the position of the study site within the country of Kenya. Inset shows the zoomed view of the sampling locality within the coastal region. The colored dots highlight the sampling locality of the study site: red= Shimoni caves (Kwale county), green=Kikambala (Kilifi county), yellow=Mavueni quarry (Kilifi county) and purple=Malindi (Kilifi county).

### Virus isolation

Samples were clarified at 4,000 x g for 10 minutes at 4°C before inoculation onto Vero E6 cells [29]. Vero E6 monolayers were prepared by seeding confluent cell monolayers in T-25cm^2^ flasks and cultivating them in MEM with antibiotics, 2 mM L-glutamine, and 10% FBS [34,35]. Then, the cultures were inoculated with 200 μL of the supernatant obtained from the bat samples. The cultivation was done for up to 14 days at 37 °C with 5% CO^2^. Cultures showing typical CPE were harvested and subjected to two blind passages to confirm viral replication [30,31]. Uninfected cells were used as negative controls. We followed virus isolation approaches previously used for bat virus isolation [32–34].

### Nucleic acid extraction, library preparation and sequencing

Viral nucleic acid extracts were made from 140 µL of supernatants using zymo research DNA/RNA kit following manufacturer’s protocol. Nucleic acids extracted were quantified using Qubit fluorometer. rRNA depletion was done using NEBNext rRNA depletion kit (Human/Mouse/Rat) (New England Biolabs, USA). First strand cDNA was generated using Superscript IV Reverse Transcriptase (Invitrogen) and random hexamer primers. The libraries were made using Illumina DNA/RNA library prep, tagmentation kit (Illumina) and index primers. Concentration and size profile of the libraries were analyzed using Agilent 2100 Bioanalyzer. Pair-end sequencing for 2×150 bases was performed using the Illumina MiSeq platform. Quality control was maintained throughout the process with negative extraction (nuclease free water processed alongside samples) and library preparation controls (no template controls). Raw sequencing data were demultiplexed and exported as FASTQ files using the MiSeq Reporter software.

### Bioinformatics

Quality control was carried out on the generated raw sequence reads in order to remove adapter sequences, short reads of less than 50 bp, and low quality bases (quality score < 30) using Trimmomatic [35]. Quality reads were aligned to the genome of host species and humans using BWA [36] for removal of host sequences. Non-host sequences were assembled using MEGAHIT [37]. Taxonomic identification of contigs was performed by the use of Kraken2 against NCBI viral reference database [38]. Contigs of viral origin were analyzed through BLASTn and BLASTx analysis against NCBI nr database [39]. Viral sequences were identified based on established thresholds, including an E-value < 1 × 10⁻⁵, amino acid or nucleotide identity greater than 60% to the closest viral relative, and query coverage exceeding 50% [40,41]. Contigs matching non-viral sequences, were excluded from downstream analysis [42].To differentiate true viral sequences from potential contamination, negative controls were sequenced and analyzed in parallel, and any viral sequences detected in these controls were subtracted from sample data [43].

### Phylogenetic analysis

Reference *UL19* sequences from the representatives of the *Alphaherpesvirinae*, *Betaherpesvirinae*, and *Gammaherpesvirinae* were retrieved from GenBank. Multiple sequence alignment was performed with Multiple Alignment with Fast Fourier Transform (MAFFT version 7) [44]. The maximum likelihood-based phylogeny estimation was carried out using IQ-TREE 2 [45,46] with 1000 replicates of ultrafast bootstraps [47]. Genetic distances (p-distance) between each pair of sequences were computed using MEGA X [48] software using nucleotide sequence of *UL19*. A heatmap was plotted using Seaborn library in Python [49] based on the distance matrix. Phylogenetic tree was visualized using Interactive Tree of Life (iTOL) v6 [50].

## Results

### Virus isolation, sequencing and assembly

Virus isolation from 170 samples (oral and rectal swabs) obtained from three species of bats (*Hipposideros* spp., *Coleura afra* and *Rousettus aegyptiacus*) was performed. All samples were inoculated onto Vero E6 cell monolayers and observed for 14 days. Cytopathic effects (CPE) were observed in two cultures of rectal samples from *R. aegyptiacus* (KIK_460_O and KIK_465_O) five days post-inoculation [31,51]. Isolates were harvested and stored at -80 °C when CPE developed in approximately 70% of the monolayer. The presence of a replicating virus was confirmed by the reproducible CPE patterns (up to 3 passages), while control wells maintained normal morphology. The two CPE positive isolates were selected for further genomic analysis.

High throughput sequencing of each of the two CPE positive isolates produced about five million paired-end reads per sample. On average 4.6 million high-quality reads per sample were retained after adapter trimming and quality filtering. De novo assembly using MEGAHIT [37] produced multiple contigs for each sample, which were assembled into partial consensus sequences of approximately 60 kb (KIK_460_O) and 70 kb (KIK_465_O). These partial genomes contain multiple alphaherpesvirus genes including *UL19*, *UL25*, and *UL30*. Based on the size of related alphaherpesviruses, the complete genome of the novel fruit bat alphaherpesvirus is estimated to be approximately 140 kb [7], meaning our partial assemblies cover approximately 43-50% of the complete genome (Supplementary File 1).

### Virus detection

#### Novel fruit bat alphaherpesvirus

Alpha herpesvirus was isolated from both CPE-positive samples (KIK_460_O and KIK_465_O). The partial consensus sequences contained the *UL19* gene (major capsid protein) in both isolates, and the *UL30* gene (DNA polymerase) in S42 (KIK_465_O). The *UL19* genes from S41 (KIK_460_O) (3,787 bp) and S42 (2,887 bp) showed 84.68% and 84.53% nucleotide identity, respectively, to the Dzifa herpesvirus reference KC701028.1. The *UL30* gene (2,846 bp) showed 86.43% identity to KC701030.1. Both isolates were genetically identical (p-distance = 0.00), confirming they represent the same viral lineage. These sequences extend the available genomic data for alphaherpesvirus, representing the first extended genomic characterization of this viral species. The phylogenetic placement of alpha herpesvirus near primate alphaherpesviruses is discussed in Supplementary Table S1.

### Phylogenetic analysis and genetic distances

#### Phylogenetic analysis of *UL19*

Phylogenetic analysis was performed using the *UL19* amino acid sequences from the Dzifa herpesvirus isolates (KIK_460_O and KIK_465_O) together with reference sequences from the *Alphaherpesvirinae* subfamily. The maximum likelihood tree placed the Dzifa herpesvirus isolates within the *Alphaherpesvirinae* subfamily (Figure 2).

**Figure 2.**
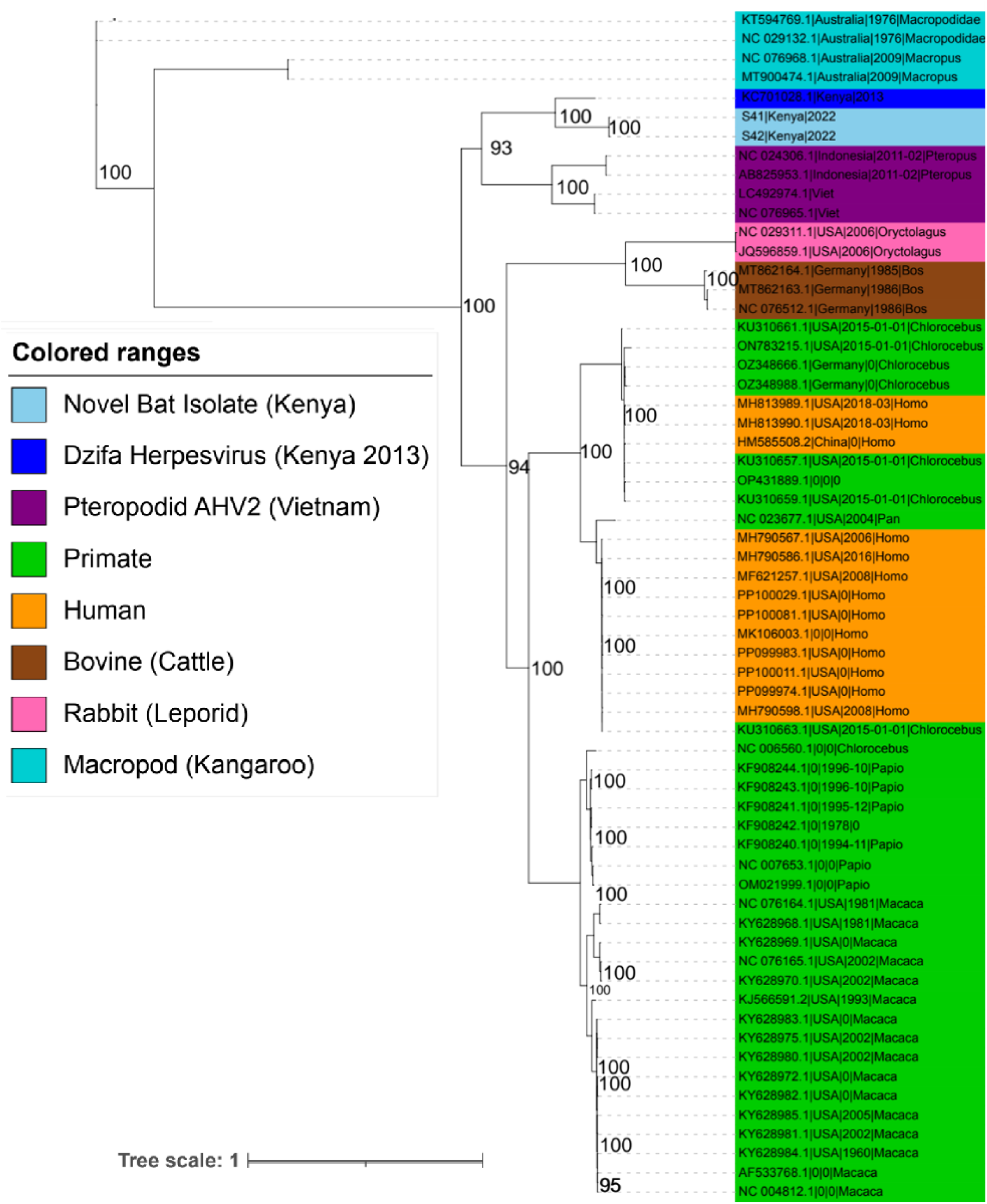
Maximum likelihood phylogenetic tree based on UL19 amino acid sequences. The tree was constructed using IQ-TREE 2 and 1,000 ultrafast bootstrap replicates. Bootstrap support values (≥70%) are shown at the nodes. Sequences from this study (S41 and S42, both Dzifa herpesvirus isolates from Kilifi, Kikambala, 2024) are shown in bold light blue. The previously described Dzifa herpesvirus isolate KC701028.1 (Dzifa herpesvirus|Kenya|2013) is shown in blue.

The alphaherpesvirus isolates from this study (KIK_460_O and KIK_465_O) clustered most closely with the previously described Dzifa herpesvirus isolate KEN709 (AHB18039.1) from Kenya (2013), with 100% bootstrap support. This clade was positioned among primate alphaherpesviruses, including *Panine alphaherpesvirus 3*, *Macacine alphaherpesvirus 3*, and *Macacine alphaherpesvirus 1*, with approximately 30% nucleotide divergence [52].

Pairwise genetic distance analysis was performed to quantify the genetic relationships among the 12 alphaherpesvirus sequences included in the phylogenetic analysis (Figure 3; Supplementary Table S2). The two isolates from this study, S41_UL19 and S42_UL19, were genetically identical (p-distance = 0.00), confirming they represent the same viral lineage. The Dzifa herpesvirus isolates ((KIK_460_O, KIK_465_O and KC701028.1) showed moderate divergence from primate alphaherpesviruses (0.23–0.28), consistent with their phylogenetic placement near primate viruses. Greater divergence was observed between Dzifa herpesvirus and pteropodid-like viruses from Asia (0.30–0.32), and between Dzifa herpesvirus and bovine/rabbit alphaherpesviruses (0.30–0.32). Specifically, S41_UL19 showed 0.245 divergence from LC492974.1_UL19 (pteropodid AHV2), 0.268 divergence from NC_023677.1_UL19 (primate), 0.300 divergence from MT862163.1_UL19 (bovine), and 0.321 divergence from JQ596859.1_UL19 (rabbit). S42_UL19 showed nearly identical distances (0.250, 0.280, 0.305, and 0.311, respectively).

**Figure 3.**
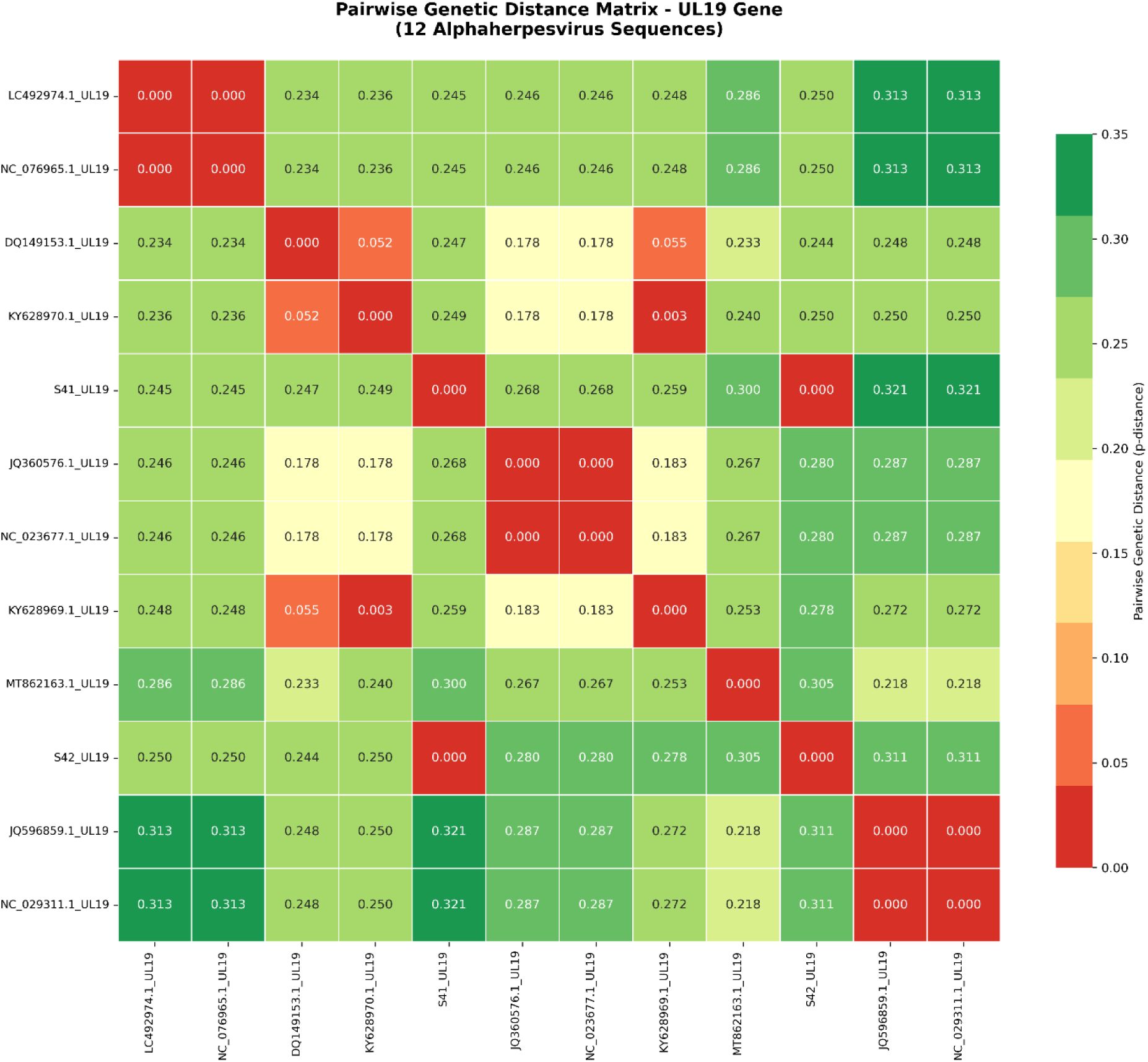
Heatmap of pairwise genetic distances (p-distance) calculated from *UL19* nucleotide sequences for 12 alphaherpesvirus isolates. Color intensity from red (low divergence) to green (high divergence) indicates genetic similarity between sequences. The distance matrix confirms that S41_UL19 and S42_UL19 are nearly identical (distance = 0.00), while Dzifa herpesvirus isolates show moderate divergence from primate alphaherpesviruses (0.23-0.28) and higher divergence from pteropodid-like viruses (0.30-0.32) and bovine/rabbit viruses (0.30-0.32). Hierarchical clustering (dendrograms) groups sequences according to host origin, consistent with the phylogenetic tree topology. The complete distance matrix is provided in Supplementary Table S2.

Hierarchical clustering of the distance matrix (Figure 3) grouped sequences according to host origin bat-derived viruses, including the pteropodid-like viruses from Vietnam and Indonesia, formed one cluster; primate viruses formed a distinct cluster; and viruses from other mammalian hosts, such as bovine, rabbit, and macropod, formed separate clusters. This host-specific clustering supports the co-evolutionary history of alphaherpesviruses with their mammalian hosts and is consistent with the phylogenetic tree topology [53].

The near-identity of S41_UL19 and S42_UL19 (p-distance = 0.00) provides strong evidence that both isolates represent the same viral strain. The moderate divergence between Dzifa herpesvirus and primate alphaherpesviruses (0.23–0.28) suggests a shared evolutionary history, while the higher divergence from pteropodid-like viruses (0.30–0.32) and bovine/rabbit viruses (0.30–0.32) indicates more distant relationships.

## 4. Discussion

This study reports the isolation and partial genomic characterization of alphaherpesvirus from Kenyan *Rousettus aegyptiacus* bats. Both isolates (KIK_460_O and KIK_465_O) were genetically identical, confirming they represent the same viral lineage. The successful isolation of alphaherpesvirus in Vero E6 cells confirms that this virus has replicative ability in mammalian cell lines, consistent with observations from previous bat herpesvirus isolations [32,34,54]. The partial consensus sequences obtained in this study (∼60 kb for S41[KIK_460_O] and ∼70 kb for S42 [KIK_465_O]) represent the first extended genomic data for alphaherpesvirus, covering approximately 43-50% of the estimated 140 kb complete genome. These sequences provide new genomic data for the *UL19* (3,787 bp in S41; 2,887 bp in S42) and *UL30* (2,846 bp in S42) genes. The nucleotide identity between alpha herpesvirus isolates and the KEN709 reference (84.53%–86.68%) indicates they belong to the same viral species but are not identical isolates, consistent with natural genetic diversity in bat herpesvirus populations [55]. The extension of genomic data is consistent with recent metagenomic studies that have similarly expanded known viral genomes through unbiased sequencing approaches [56,57].

The phylogenetic analysis (Figure 2) and pairwise genetic distance analysis (Figure 3; Supplementary Table S2) provide complementary evidence for the evolutionary relationships of the viruses identified in this study. S41_UL19 and S42_UL19 were genetically identical (p-distance = 0.00), confirming they represent the same viral lineage. This near-identity, combined with their co-isolation from the same bat species (*R. aegyptiacus*) from the same location (Kikambala, Kilifi County), suggests they were derived from a common source or represent a single circulating strain [10]. The moderate divergence between Dzifa herpesvirus and primate alphaherpesviruses (0.23-0.28) is consistent with their phylogenetic placement within the *Alphaherpesvirinae* subfamily near primate viruses. The higher divergence from alphaherpesvirus-like viruses (0.30-0.32) and bovine/rabbit viruses (0.30-0.32) indicates that alphaherpesvirus is more distantly related to these lineages [58].

The genetic distance analysis further revealed that the alphaherpesvirus isolates (S41, S42, [KIK_460_O and KIK_460_O] and KC701028.1) form a distinct cluster, with approximately 0.30 divergence from pteropodid-like viruses. This level of divergence is consistent with genus-level differences within the *Alphaherpesvirinae*, suggesting that alphaherpevirus represents a distinct viral species within this subfamily [3]. The clustering of sequences by host origin in both the phylogenetic tree and the heatmap provides strong evidence for host-specific evolution of these alphaherpesvirus lineages [59].

The detection of alphaherpesviruses in *Rousettus aegyptiacus*, which roosts in caves, indicates ecological interfaces where viral exchange can occur [60]. Human activities in coastal Kenya, including quarrying, tourism, and deforestation, are increasing contact with bat habitats, and this increases the risk of exposing people to bat-borne viruses [61,62]. The Shimoni cave system is a tourist site where visitors have close contact with bat roosts and could act as intermediate hosts. Although herpesviruses are mostly host-specific, there are some exceptions, including *M. alphaherpesvirus 1* (B virus), which shows that the result of cross-species transmission can result in severe disease [4]. The phylogenetic closeness of alphaherpesvirus to primate alphaherpesviruses raises questions about its potential to infect other hosts other than bats, but there is no direct evidence of this spillover currently [59]. Future research should focus more on serological surveys on human populations residing near bat roosts to determine exposure and possible asymptomatic infection [61].

## 5. Conclusion

This study reports the isolation and partial genomic characterization of alphaherpevirus from Kenyan *Rousettus aegyptiacus* bats. The partial consensus sequences obtained (∼60 kb for S41 and ∼70 kb for S42) represent the first extended genomic data for this virus, covering approximately 43-50% of the estimated 140 kb complete genome. Phylogenetic analysis confirmed that Dzifa herpesvirus clusters within the *Alphaherpesvirinae* subfamily near primate viruses, while pairwise genetic distance analysis revealed that S41_UL19 and S42_UL19 are genetically identical (p-distance = 0.00), confirming they represent the same viral lineage. The moderate divergence from primate alphaherpesviruses (0.23-0.28) and higher divergence from other alphaherpesvirus lineages (0.30-0.32) provides quantitative evidence for the evolutionary relationships of these viruses. The capacity of alphaherpesvirus to replicate in Vero E6 cells highlights the viral diversity at the bat-primate-human interface in coastal Kenya. Serological studies are required to determine whether alphaherpesvirus circulates exclusively in bats or has already crossed species boundaries, as has been the case in other bat-borne viruses with the potential to be zoonotic.

## Declaration

### Ethics approval and consent to participate

Scientific and ethics approval was given by the Scientific and Ethics Review Unit (SERU) of the Kenya Medical Research Institute (KEMRI) under Protocol No. KEMRI/SERU/CVR/002/4729. The National Commission of Science, Technology, and Innovation (NACOSTI) issued a research license (No. NACOSTI/P/23/27916). All procedures followed institutional guidelines for wildlife research. Consent for participation was not applicable as the study involved only animal subjects.

### Consent for publication

Not applicable

### Availability of data and supporting materials

The datasets generated and/or analysed during the current study are not publicly available due to ongoing analyses and future publications but are available from the corresponding author on reasonable request. The assembled consensus viral genome sequences are provided in Supplementary File 1. The pairwise genetic distance matrix is provided as Supplementary Table S2.

### Authors’ contributions

GK: Conceptualization, investigation, writing original draft, visualization. JB: Supervision, writing, review and editing. JK: Supervision, writing, review and editing. SL: Methodology, data curation. JL: Resources, supervision, writing, review and editing. FE: Funding acquisition, project administration, supervision, writing, review and editing. All authors read and approved the final manuscript.

## Acknowledgements

We are grateful for the technical and logistical support of the field teams of the Department of Entomology, Kisian, Kisumu.

## Supplementary Materials

The following supplementary materials are available.

**Supplementary Table S1. Summary of viral sequences identified in bat samples S41 and S42.**

Detailed information for viral contigs identified from the two CPE-positive samples (S41 and S42). Columns include: sample ID, consensus genome, gene or viral protein, top BLAST hit with GenBank accession, query coverage (%), nucleotide or amino acid identity (%), E-value, and assigned virus lineage.

**Supplementary Table S2**. Pairwise genetic distances (p-distance) calculated from *UL19* nucleotide sequences for 12 alphaherpesvirus isolates. Values represent the proportion of nucleotide differences between sequences. S41_UL19 and S42_UL19 show 0.000 distance, indicating they are nearly identical isolates. The table includes bat-associated viruses (LC492974.1, NC_076965.1, S41_UL19, S42_UL19), primate viruses (DQ149153.1, KY628970.1, JQ360576.1, NC_023677.1, KY628969.1), bovine virus (MT862163.1), and rabbit viruses (JQ596859.1, NC_029311.1).

**Supplementary File 1. Assembled viral contig sequences in FASTA format.**

Contains the two consensus viral genome sequences assembled from samples S41 and S42 (KIK_460_O and KIK_465_O). Each consensus genome contains multiple alphaherpesvirus genes identified in this study. Sequence headers indicate sample ID and genome description. Individual contig sequences used for assembly are also included for transparency.

